# A rapid, cost-effective, efficient method for total RNA extraction from pure bacterial cultures and complex microbiomes and its effectiveness in deciphering functionally active microbial populations

**DOI:** 10.64898/2025.12.02.691598

**Authors:** Abhinaba Chakraborty, Bomba Dam

## Abstract

Extraction of high-quality RNA from microbial samples, especially complex microbiomes like the gut, remains a significant challenge. The instability of RNA, coupled with the abundance of RNases and difficult-to-lyse microbial cells, often results in degraded or low-yield RNA, hampering downstream applications. To overcome this critical bottleneck, we developed a simple, rapid, cost-effective, and efficient microbial RNA extraction method, which is equally effective for pure cultures and complex microbiomes. The method utilises low pH (4.7) and high temperature to lyse the cells, followed by rapid cooling to induce misfolded renaturation of genomic DNA and proteins. During the phase separation step, RNA remains in the aqueous phase and is subsequently precipitated using chilled isopropanol. The RNA suspended in double-distilled water (pH 6.5) remained stable for six months at −20 °C. Further, we compared amplicon microbiome profiles generated from metagenomic DNA and cDNA (synthesised from extracted meta-RNA) derived from the broiler gut, taken as a model system. Our findings reveal that the cDNA-based amplicon microbiome composition more accurately deciphers the metabolically active population in gut environments that aligns with culture-based enumeration of viable microbes. This proof-of-concept study underscores the importance of meta-RNA approaches for functional microbiome analysis and provides a reliable RNA isolation method that can be broadly adopted for studies aiming to resolve active microbial community structure.

## 1.0 Introduction

The extraction of high-quality, intact RNA is a vital step in understanding gene expression and cellular function (Fleige & Pfaffl, 2006). However, RNA is inherently unstable and more chemically reactive than DNA, making its extraction particularly challenging (Thatcher, 2015). Unlike DNA, RNA is single-stranded and possesses a hydroxyl (-OH) group at the 2’ position of its ribose sugar, which makes RNA more prone to hydrolysis (Fang, Xiao, Jun, Onishi, & Kool, 2023; Harp et al., 2022). These properties, coupled with the ubiquitous presence of ribonucleases, complicate the process of RNA extraction across biological systems. Even trace amounts of RNases, which are extremely stable and can remain active under harsh chemical conditions, are enough to significantly degrade RNA during the isolation process. In addition to chemical instability, RNA is highly sensitive to environmental conditions such as temperature, pH, and ionic strength. Fluctuations in any of these parameters can trigger rapid degradation (Kornienko, Aramova, Tishchenko, Rudoy, & Chikindas, 2024). Microbial RNA, from bacteria and archaea, presents a unique set of challenges that are not typically encountered with eukaryotic RNA (Hör, Gorski, & Vogel, 2018). First, bacterial cells often have complex and resilient cell walls, particularly in Gram-positive species, which make cell lysis more difficult and inconsistent (Danaeifar, 2022). Insufficient lysis directly affects RNA yield and quality. Secondly, microbial RNA has an extremely short half-life. In many bacterial species, mRNA degrades within minutes due to the high activity of endogenous RNases, making it critical to stabilise RNA immediately upon cell harvest (Houseley & Tollervey, 2009; Mohanty & Kushner, 2016). Consequently, RNA extraction procedures must be performed quickly and under controlled conditions to preserve RNA integrity. Even then, contamination with genomic DNA, inefficient cell lysis, and incomplete removal of proteins or phenol residues can further impair RNA quality. These general difficulties are amplified when working with low biomass or hard-to-lyse cells, factors that frequently occur in environmental and clinical studies.

These challenges become even more pronounced when dealing with complex microbial communities or “microbiomes,” particularly those from environmental or host-associated niches such as the gut. Microbiomes are typically composed of hundreds to thousands of microbial species, varying widely in abundance, physiology, and lysis susceptibility (Berg et al., 2020). The co-extraction of host RNA and the presence of hard-to-remove contaminants like organic acids, bile salts, or polysaccharides from gut samples can compromise RNA purity and inhibit downstream reactions such as reverse transcription. Despite these difficulties, RNA-based approaches seem attractive in microbiome research due to their ability to provide a snapshot of the “active” members of the community, which is not possible using DNA-based conventional metagenomics (Wang, Zhang, Liu, Zhang, & Sun, 2021; Yang et al., 2022). This discrepancy is critical in environments such as the gut, where a significant fraction of the microbial DNA pool (up to 30% in some studies) may originate from dead cells (Bellali, Lagier, Raoult, & Bou Khalil, 2019; Fu et al., 2018). The presence of relic DNA can obscure community dynamics and confound the interpretation of treatment effects, disease associations, or environmental responses.

However, the RNA isolation step remains the most significant technical bottleneck in microbiome studies. In this study, we developed a simple, efficient, and cost-effective method for extracting microbial RNA from both pure bacterial cultures and complex gut environments. The method was rigorously quality-checked and validated for its applicability across multiple downstream analyses, including qPCR, and high-throughput 16S rRNA gene amplicon sequencing. Notably, it proved highly effective in capturing the transcriptionally active fraction of gut microbial communities, offering a reliable alternative to conventional protocols, particularly in settings where accessibility, affordability, and reproducibility are critical.

## 2.0 Materials and Methods

### 2.1 Bacterial cultures, reagents and consumables

Two Gram-positive (*Staphylococcus aureus* MTCC 96, *Bacillus subtilis* MTCC 739) and two Gram-negative bacteria (*Escherichia coli* MTCC 739, *Pseudomonas aeruginosa* MTCC 3301) were used for RNA isolation. Additionally, cecal and intestinal contents of 42-day-old broiler birds were used for evaluation of the method. The reagents used for this study are: extraction solution (50 mM glucose, 10 mM EDTA, adjusted to pH 4.7 using acetic acid, autoclaved and refrigerated); 1X TAE buffer (40 mM Tris, 1 mM EDTA, pH adjusted to 7.0 using acetic acid, autoclaved); water saturated phenol (pH 4.7); chilled isopropanol; and chilled ethanol (70%). All plasticwares, like microcentrifuge tubes, micropipettes and microtips, were autoclaved before use.

### 2.2 RNA extraction procedure

Total RNA from pure cultures and gut samples were extracted following the procedure described in Figure 1a. For this, 2 ml of cell mass from active log-phasic cultures (OD_600_ ∼ 0.4-0.6) of pure bacteria was suspended in 700 µl of extraction solution. For cecal and intestinal meta-RNA preparation, 20 mg of fresh contents were taken and suspended in the extraction solution. To these suspensions, 600 µl of water-saturated phenol (pH 4.7) was added. The mixtures were then briefly vortexed and subjected to heat treatment (95°C for 4 minutes) in a dry/water bath. Complete lysis of bacteria was indicated by the formation of a transparent solution. Following lysis, the tubes were placed on ice for 5 minutes, leading to the formation of a white precipitate. To this mixture, 600 µl of chilled chloroform was added and mixed gently without vortexing. Then, the tubes were centrifuged at 12,000 g for 10 minutes at 4°C, with the deceleration rate set to 0, to avoid phase mixing. The upper aqueous layers were carefully transferred to new tubes, and 0.7 volume of chilled isopropanol was added and mixed properly. The mixtures were kept on ice for 5 minutes to facilitate RNA precipitation. The tubes were then centrifuged at 12,000 rpm for 10 minutes at 4°C. The RNA pellets were subsequently washed by adding 70% (v/v) chilled ethanol. The tubes were subjected to a final centrifugation at 12,000 rpm for 10 minutes at 4°C. Following this, the 70% ethanol solutions were discarded, and the RNA pellets were air-dried for 10 minutes. Finally, the dried pellets were re-suspended in 50 µl autoclaved double-distilled water for subsequent analyses.

**Figure 1:**
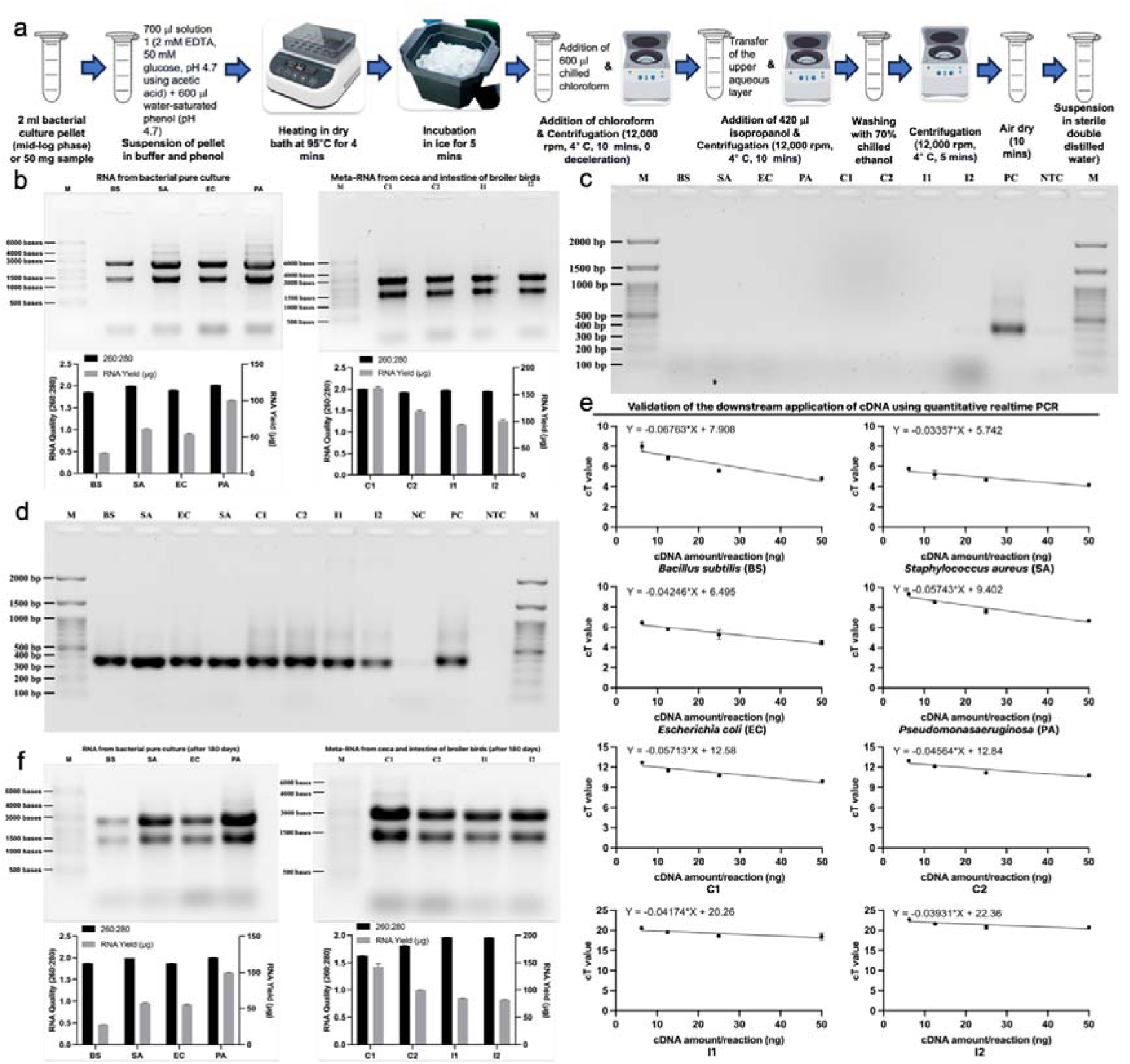
Extraction of microbial RNA and their validation. **a**, Flowchart of RNA extraction process. b, Gel image showing microbial RNA from pure cultures and gut samples. Bar diagrams below the gel images show total RNA yield and 260:280 ratio of the extracted RNA. c, Gel image showing negative PCR amplification on RNA samples. d, Gel image showing amplification of V1-V2 region of 16S rRNA region performed on cDNA, synthesized from the RNA samples. e, Linearity trend of cDNA dilutions through quantitative real-time PCR shows downstream applicability of the cDNA. f, Gel image showing stability of RNA samples after 6 months of storage at −20°C. The bar graphs below each gel show RNA yield and 260:280 ratio. All experiments were performed in triplicates and mean values were plotted with error bars showing standard deviation BS= *Bacillus subtilis*, SA= *Staphylococcus aureus*, EC=*Escherichia coli*, PA= *Pseudomonas aeruginosa*, I1 and I2= Intestine content 1 and 2, C1 and C2= Cecal content 1 and 2, NC= Negative Control, PC= Positive Control, NTC= No Template Control, M= DNA or RNA marker

### 2.3 Determination of RNA quality, quantity and stability

Agarose gel electrophoresis was employed to assess RNA integrity. A 1.5% agarose gel was prepared using 1X TAE buffer, pH 7.0 (adjusted using acetic acid). The same buffer was also used for running the gel. For visualisation, a higher ethidium bromide concentration (0.5 µg/ml) was used. The gel was run at 100 V until the dye front had migrated half the gel, and RNA bands were visualised using a gel documentation system (GelDoc, Biorad, USA).

RNA concentration and purity were measured using a NanoDrop (Epoch2, Biotek, USA). Absorbance was measured at 260 nm (A_260_) to determine RNA concentration, and the ratio of A_260_: A_280_ was used to assess RNA purity. The RNA concentration was calculated using the formula: RNA concentration (ng/µL) = A_260_×dilution factor×40.

The RNA samples were kept at −20°C for 6 months and checked again on agarose gel and Nanodrop spectrophotometer (Biotek, USA).

### 2.4 Comparison with other commercially available kits and methods

To check effectiveness, total RNA was isolated from the same pure cultures and gut samples, using the developed method, and the yield and quality were compared with other commercially available kits and reagents. For this, the RNeasy mini kit (Qiagen, USA) and the Trizol reagent (Thermo Fisher Scientific, USA) were used.

We have also made a framework-based schematic comparison of our RNA extraction method with other developed methods from their literatures. For this, we did a search in Web of Science with this prompt: ((TS=(RNA extraction)) AND TS=(Bacteria)) AND TS=(Method), which gave a total of 638 results. We read all the articles and selected only those where an indigenous RNA extraction method was developed (Table 2).

### 2.5 Detection of DNA contamination in RNA samples using end-point PCR

To verify the absence of contaminating DNA in the RNA samples, PCR amplification of the V1-V2 region of the 16S rRNA gene was performed. Each reaction mixture contained the following components: 2.5 µl of 10x PCR buffer, 0.5 µl of 10 mM dNTP (abclonal, USA), 0.5 µl of forward primer (8F: AGAGTTTGATCITGGCTCAG: 10 µM), 0.5 µl of reverse primer (338R: CTGCTGCCTCCCGTAGG: 10 µM), 100 ng of template RNA sample, 0.125 µl *Taq* DNA polymerase (abclonal, USA) and sterile water to bring the final volume to 25 µl. For the positive control, 100 ng of *Escherichia coli* genomic DNA was used as the template. For the no-template control, 2 µl of RNase-free water was added in place of the template.

The thermal cycler was programmed with the following conditions: initial denaturation at 95°C for 5 minutes, followed by 30 cycles of denaturation at 95°C for 30 seconds, annealing at 58°C for 45 seconds, and extension at 68°C for 1 minute. A final extension step at 68°C for 5 minutes completed the PCR amplification. After the PCR, the amplification products were visualised using agarose gel electrophoresis.

### 2.6 cDNA synthesis from total RNA

The synthesis of cDNA from total RNA was conducted using the iScript cDNA Synthesis Kit (Bio-Rad, USA). A 20 µl reaction includes 4 µl of 5X iScript Reaction Mix, 1 µL of iScript Reverse Transcriptase, up to 1 µg of total RNA, and nuclease-free water to reach the final volume of 20 µl. Following thorough mixing and brief centrifugation to consolidate all components, the reaction tube was incubated in a thermal cycler (Biorad, USA) under specific conditions: initial priming at 25°C for 5 minutes, followed by reverse transcription at 46°C for 20 minutes, and final inactivation of reverse transcriptase at 95°C for 1 minute.

### 2.7 End-point and real-time quantitative PCR validation of cDNA

The synthesised cDNAs were validated using both end-point and real-time quantitative PCR. For end-point PCR validation, the reactions were run in a similar manner as mentioned previously. For real-time quantitative PCR (qPCR), the cDNA samples were serially diluted with sterile water. The dilutions were used for amplification of the V1-V2 region of the 16S rRNA gene. The qPCR reactions were performed using Genious 2X SYBR Green Fast qPCR mix (abclonal, USA). Each reaction mixture had a final volume of 20 µL, consisting of 10 µL of Genious 2X SYBR Green Fast qPCR Mix, 0.4 µl of each primers (10 µM), and 2 µL of the cDNA template dilutions, 0.4 µl ROX reference dye II, with the remaining volume made up with sterile water. The following thermal cycling conditions were used: an initial hold stage at 50°C for 2 minutes and 95°C for 10 minutes, followed by a PCR stage of 40 cycles of denaturation at 95°C for 15 seconds, amplification at 60°C for 1 minute. Melting curve analysis was subsequently performed to confirm the specificity of the amplification. The real-time qPCR was conducted in triplicate for each cDNA dilution to ensure reproducibility and reliability of the results. Cycle threshold (Ct) values were recorded for each reaction, and the data were analysed to assess the efficiency of amplification and the linearity across different dilutions.

### 2.8 16S rRNA gene amplicon library generation and sequencing

Cecal and intestinal contents of three broiler birds were pooled for the extraction of metagenomic DNA and meta-RNA. Metagenomic DNA were extracted using SPINeasy DNA Kit for Feces/Soil (MP Biomedicals, Santa Ana, CA), following the manufacturer’s instructions. From the same lot of samples, meta-RNA was extracted using our established protocol, checked for DNA contamination and used for cDNA synthesis. 16S rRNA (V1-V2 region) amplicon libraries were generated (in triplicate) on both the metagenomic DNA and cDNA using barcoded primers, gel eluted, and sequenced on an Ion PGM 318 chip with 400-base read length chemistry (Banerjee et al., 2018).

### 2.9 Bioinformatic analysis

Initial quality assessment of the raw sequences was performed using FastQC. Raw reads were then trimmed using Cutadapt for removing low-quality bases and sequences outside the length range of 100-370 bp with quality thresholds set to Q30 for leading and Q20 for the trailing bases. Trimmed fastq files were organized and imported to QIIME2 using a single-end PHRED-33 manifest file format (Hall & Beiko, 2018). Denoising and further quality filtering of the imported sequences were performed using DADA2 plugin in QIIME2. Denoised outputs included a table of feature frequencies and representative sequences. Taxonomic classification of representative sequences was performed using a pre-trained Silva 138 classifier targeting the V1-V2 region of the 16S rRNA gene (Quast et al., 2012). Taxonomic abundances at levels 2 to 4 were calculated using the Taxa collapse plugin in QIIME2. Alpha diversity indices, including Shannon, Simpson, and observed features, were calculated using QIIME2 plugins. Rarefaction curves were generated to assess sequencing depth and richness using a range of metrics (e.g., Shannon, Simpson) with a stepwise depth increment from 1,000 to 30,000 sequences. Beta diversity was computed with a rarefaction depth of 30,000 sequences.

Predictive functional profiling of microbial communities was performed using PICRUSt (Phylogenetic Investigation of Communities by Reconstruction of Unobserved States) (Douglas, Beiko, & Langille, 2018). For this, a closed-reference OTU picking approach was employed to ensure compatibility with the PICRUSt database. Functional metagenomic predictions were made using the normalised OTU table. Predicted functional profiles were analysed to identify differences in metabolic pathways between sample groups. Statistical comparisons were performed using STAMP (STatistical Analysis of Metagenomic Profiles).

### 2.10 Culture-based analysis

For culture-based analysis, twelve selective and differential media were used for enumerating different members of the phylum Bacteroidota, Firmicutes and Proteobacteria (Supporting information Table 1). The cecal and intestinal contents used for metagenomic DNA and meta-RNA preparation were serially diluted, and 0.1 ml of appropriate dilutions were spread on different media plates. The plates were incubated at 37°C either aerobically or anaerobically, based on the requirements. After incubation for the desired period, the total CFU count was done, and CFU per gram of cecal and intestinal contents were determined. The media used for selectively enumerating Firmicutes members include aerobically incubated plates of Hicrome M-Enterococci agar, Mannitol salt agar, bile esculin agar supplemented with kanamycin (20 g ml^-1^) and sodium azide (0.15 mg ml^-1^), and anaerobically maintained plates of *Lactobacillus* MRS agar and reinforced clostridial agar. Bacteroidota members were selectively enumerated on anaerobically maintained plates of bile esculin agar supplements with vitamin K1 (0.01 mg ml^-1^), haemin (0.01 mg ml^-1^), and kanamycin (50 μg ml^-1^); aerobic incubation of plates of nutrient agar with kanamycin (50 μg ml^-1^); and HP6 agar with glucose (50 mg ml^-1^). For enumeration of proteobacteria members, Hicrome *E. coli* agar, Hicrome *Klebsiella* agar, Hicrome *Vibrio* agar, and *Salmonella-Shigella* agar. For the phylum Campylobacterota, *Campylobacter* agar plates were aerobically incubated (Banerjee et al., 2018).

**Table 1:**
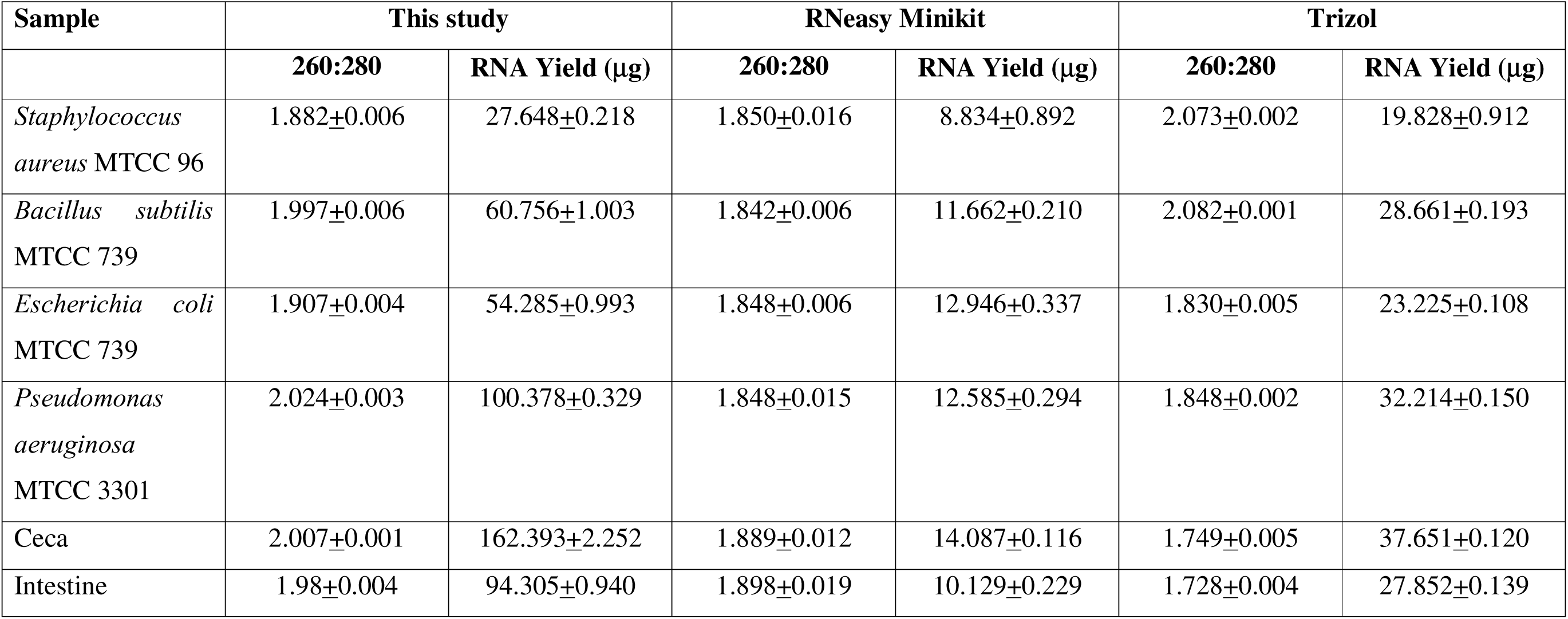
Comparison of the RNA extraction method (this study) with other commercially available kits. Data represented as mean ± standard deviation.

**Table 2:**
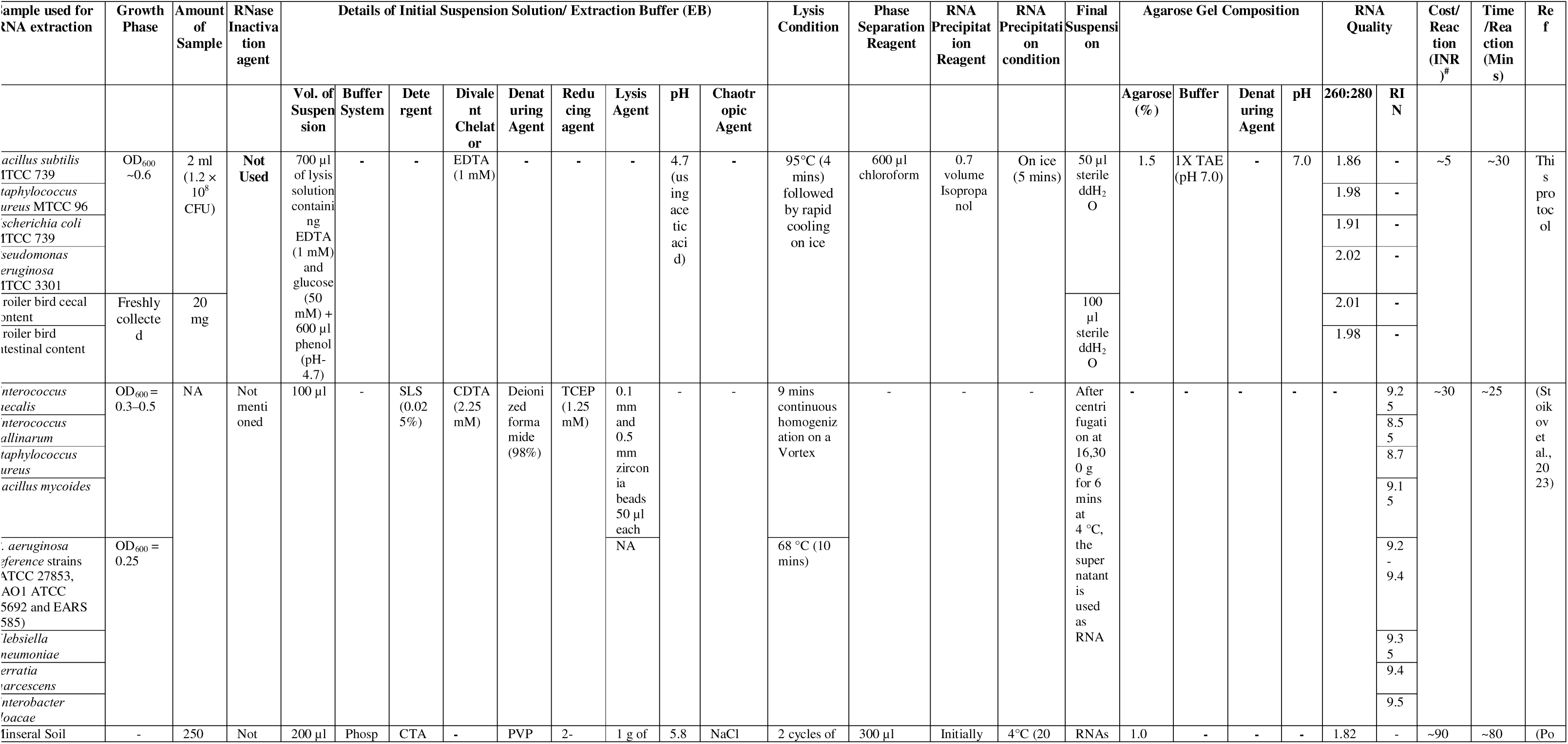
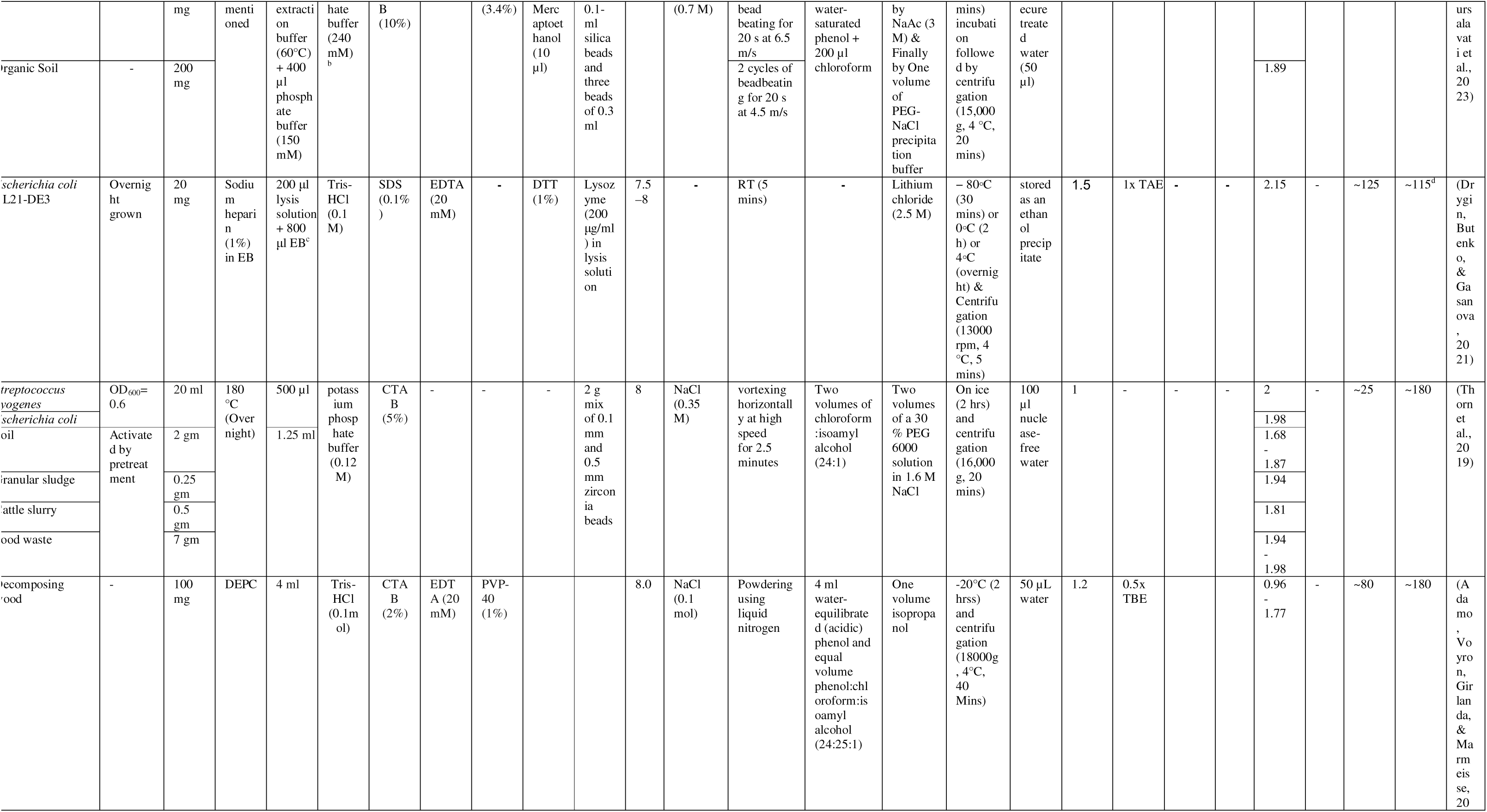
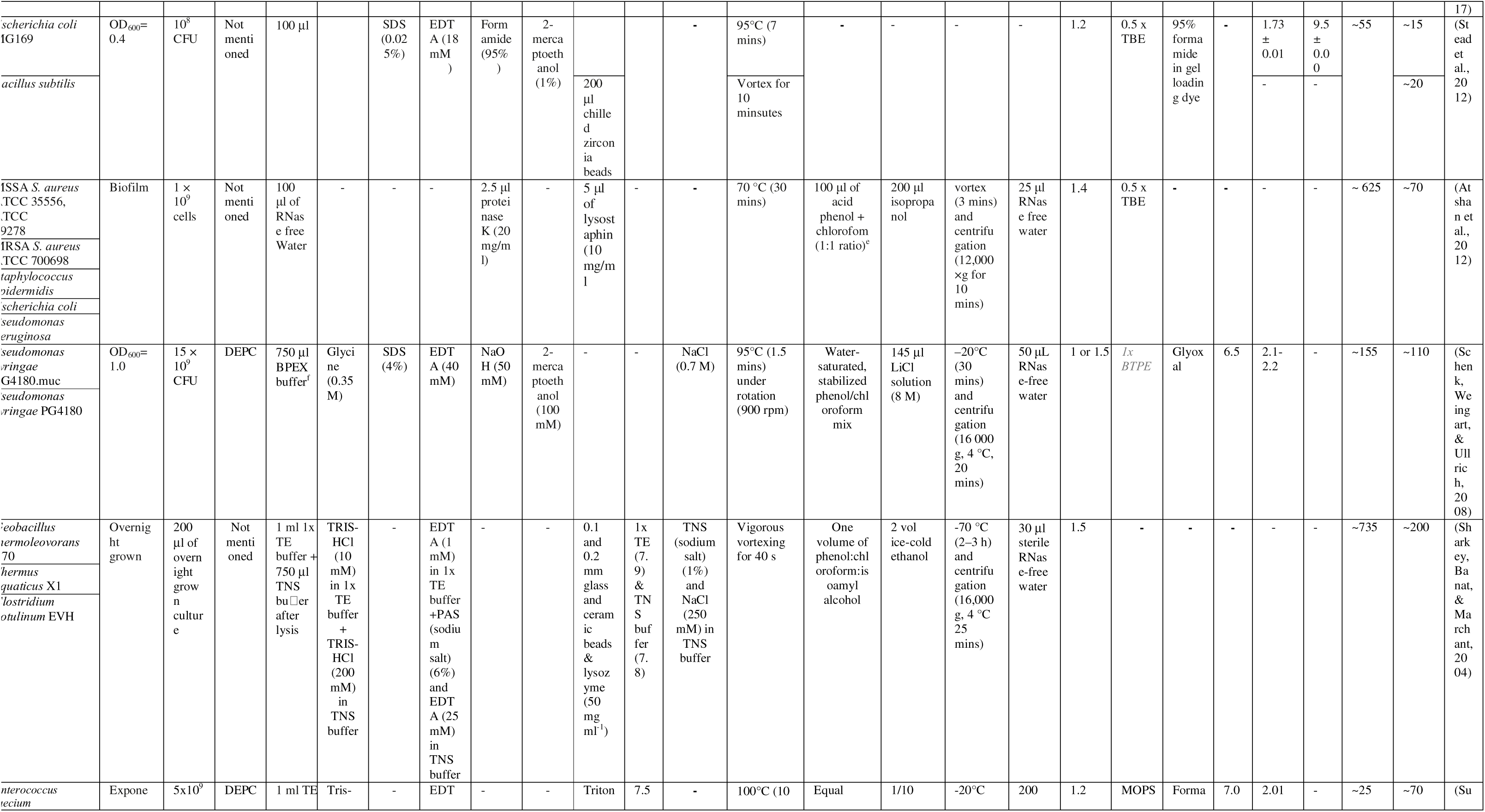
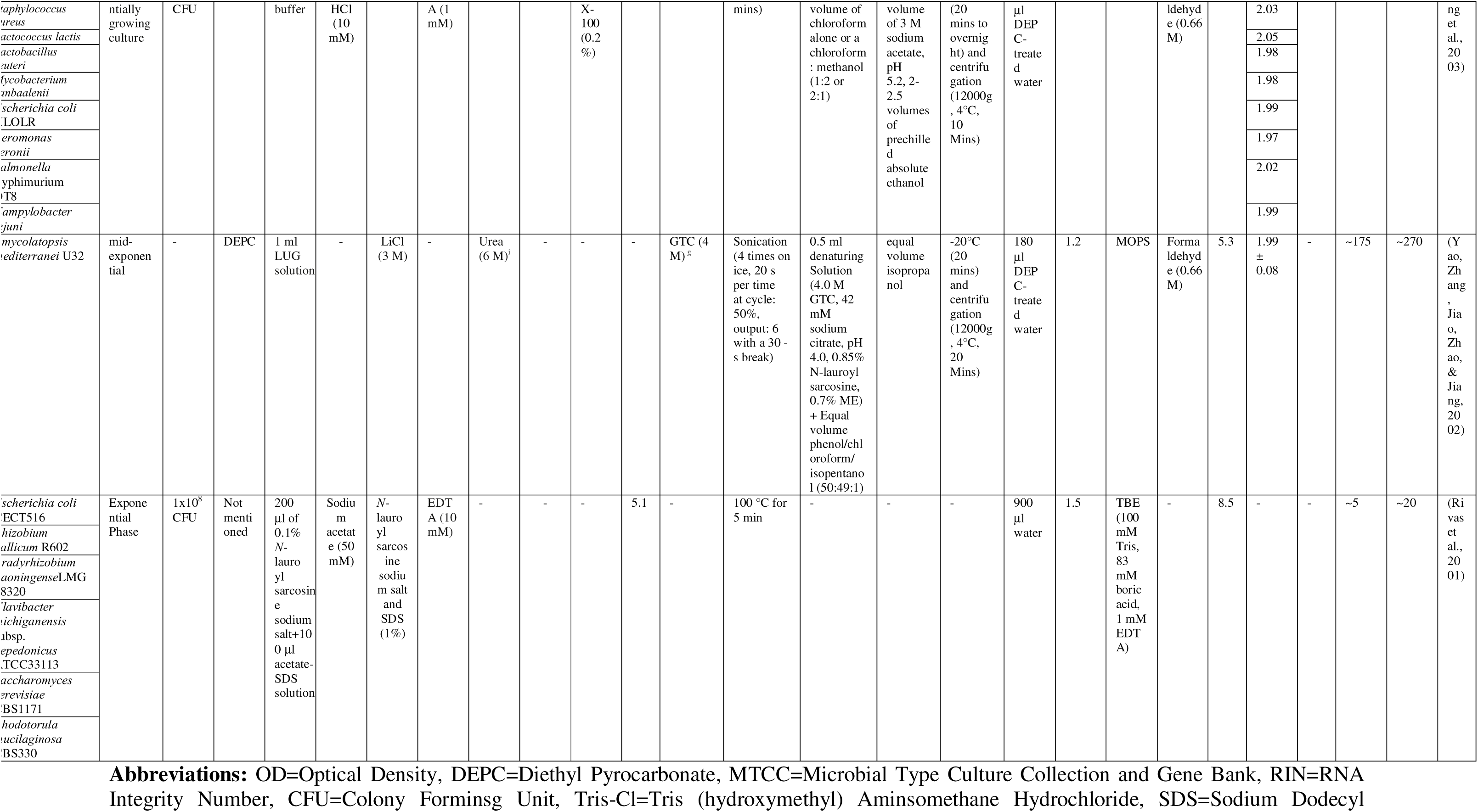
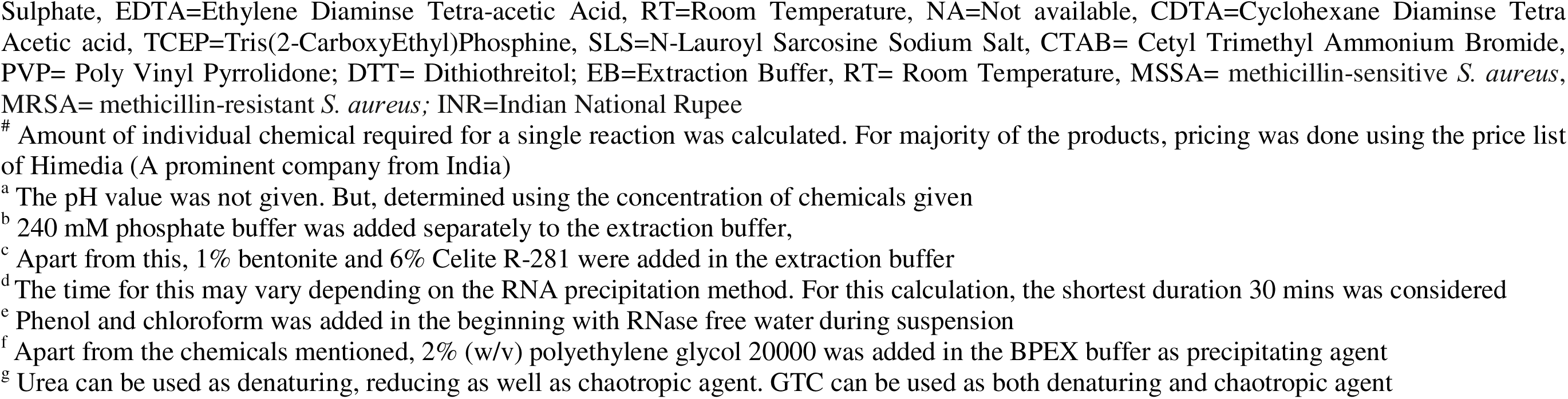
Comparison of the developed RNA extraction process with other methods. A search with the prompt: “((TS=(RNA extraction)) AND TS=(Bacteria)) AND TS=(Method)” was done in web of science. After screening 638 results, those were selected for comparison, where the indigenous RNA extraction method was developed in the last 25 years.

### 2.11 Figure preparation

The figures used in this study were prepared using GraphPad Prism and STAMP (STastistical Analysis of Metagenomic Profiles).

## 3.0 Results

### 3.1 Successful isolation of DNA-free RNA and downstream application

An efficient method for extracting meta-RNA from pure cultures and complex microbiome samples was developed and evaluated (Figure 1a). Initially, RNA was isolated from *Bacillus subtilis*, *Staphylococcus aureus*, *Escherichia coli*, and *Pseudomonas aeruginosa* with a 260:280 ratio of 1.8-2.0. Additionally, total RNA yield was very high (27-100 μg from approximately 2 × 10^8^ cells), suggesting the suitability of the RNA isolation protocol from both Gram-positive and negative bacteria (Figure 1b). Then, the method was employed for the extraction of meta-RNA from cecal and intestinal contents of broiler birds. Again, a high RNA yield with a 260:280 ratio of ∼2.0 was observed. In agarose gel, three distinct bands corresponding to 23S, 16S and 5S rRNA genes were observed for all the samples (Fig. 1b). No bands near the wells were observed, suggesting the absence of DNA contamination. Further, the absence of genomic DNA in the extracted RNA was confirmed by a negative PCR with the 16S rRNA gene (V1-V2 region)-specific primers (Fig. 1c). However, the same PCR reaction yielded positive products when cDNA (synthesized from the total RNA) was used as template (Fig. 1d). The suitability of the cDNA in real-time quantitative PCR was also evaluated using the same 16S rRNA gene-specific primers. The Ct values obtained, with increasing concentration of cDNA yield linear curves, signifying their competence for downstream analyses (Fig. 1e). Interestingly, the RNA samples remained stable in EDTA-free distilled water for 6 months at −20°C (Fig. 1f). Compared to the Trizol method (Thermo Fisher Scientific, USA) and the RNeasy kit (Qiagen, USA), our protocol yielded higher quality and greater quantities of total RNA across all six samples (Table 1). This method is easy, rapid, cost-effective, and efficient compared to the other available methods (Table 2).

### 3.2 Culture-dependent and -independent analysis of gut microbiomes

To assess the reliability of RNA-based microbiome profiling in identifying active microbial populations, we conducted both culture-dependent and culture-independent analyses of the cecal and intestinal contents of broiler birds. In both DNA and RNA-based approaches, the microbiomes of the ceca and the intestine differed significantly (Supporting information Figure S1a, S1b). Even in individual gut fractions, the microbiome profiles obtained by the two culture-independent methods significantly varied at both phylum and class levels (Supporting information Figure S1c, S1d). The rarefaction curves for all samples reached a plateau with increasing depth. Although the Shannon and Simpson indices of Ceca DNA- and RNA-based data are almost the same, both the values for the intestine are significantly higher in the cDNA-derived metagenome, indicating a more diverse active intestinal microbiome (Figure 2b). A Similar trend was observed in the weighted unifrac distance matrix (Figure 2c). The number of observed features (earlier in QIIME1 referred to as OTU or Operational Taxonomic Unit), respectively, for the DNA- and RNA-based microbiomes of the ceca were 439 and 814; and the intestine were 265 and 309. The relative abundance of Firmicutes in ceca was 78.57% and 52.53% in DNA and RNA-based microbiomes, with Clostridia (72.54% and 50.93% in DNA and RNA-based microbiomes, respectively) being the dominant class in both cases (Figure 2a). A complete opposite phenomenon was observed in intestinal samples, where the relative abundance of Firmicutes in DNA- and RNA-based microbiomes was 3.91% and 44.03% respectively, with the primary class being Bacilli (found in RNA-based microbiome). Notably, the culture-based enumerations align with the RNA-based microbiome data, with total Firmicutes load being 1.6 x 10^12^ and 7.2 x 10^10^ CFU g^-1^, respectively, in the ceca and intestine (Figure 2e). Even in reinforced clostridial agar, the CFU g^-1^ count was higher in the ceca (1.4 × 10^12^) than in the intestine (6.7 x 10^8^) (Supporting information Figure S5). Again, in sync with the RNA-based microbiome, lactic acid bacterial load in MRS agar was higher in the intestine (6.3 × 10^10^ CFU g^-1^) than in the ceca (1.2 × 10^10^ CFU g^-1^). Similarly, Bacteroidota (earlier Bacteroidetes) has a notably higher representation in RNA-based cecal microbiome (31.45%) compared to the DNA-based one (8.21%), with Bacteroidia being the prominent class (Figure 2a). However, the phyla are poorly represented in the intestinal microbiomes. In the culture-based enumerations, the total Bacteroidota counts were 1.7 x 10^12^ and 1.1 x 10^9^ CFU g^-1^ in cecal and intestinal contents, respectively (Figure 2e). Bile esculin agar, supplemented with vitamin K1, hemin and kanamycin, is selective for the genus *Bacteroides*. Its load in the ceca and intestine were 6.0 x 10^11^ and 6.8 × 10^8^ CFU g^-1^, respectively (Supporting information Figure S5). The prominent marker of gut health, i.e., Firmicutes: Bacteroidota ratio derived from the RNA-based microbiome, was 1.67 and 112.33 in the ceca and intestine, respectively. These values correspond well to the culture-based ratios (1.38 and 69.19), but the DNA-based values were completely different for the gut segments (9.56 and 1.95 for ceca and intestine, respectively) (Figure 2f). The Proteobacteria are found to be abundant in the intestine in both DNA and RNA-based microbiomes. Similar to the Firmicutes data in ceca, both Proteobacteria and its abundant class Gammaproteobacteria are overrepresented (almost double) in the DNA-based microbiome (∼86%) than the RNA-based one (∼47%). A comparable trend was noted for other phyla in DNA and RNA-based microbiomes (Figure 2e, Supporting information Figure S2, S3, S5). As Firmicutes and Proteobacteria are the two prominent phyla in the intestine, a new gut health indicator ratio was used in this study, i.e., Firmicutes: Proteobacteria, that gave consistent values in both RNA-based microbiomes and the culture-based enumerations (Figure 2g).

**Figure 2:**
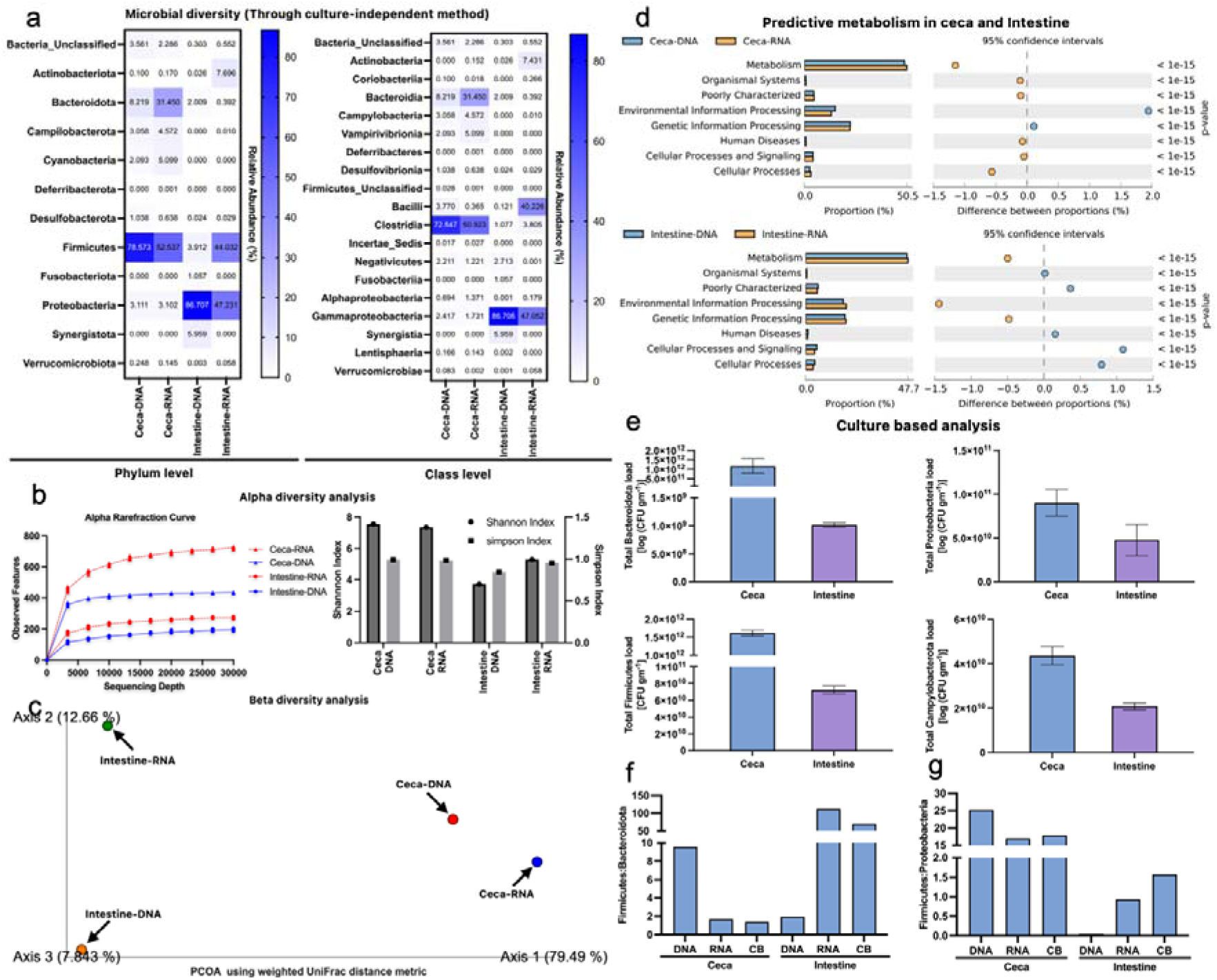
RNA based microbiome represent active microbial population. **a**, Culture-independent microbial diversity analysis of ceca and intestine of broiler using metagenomic DNA and meta-RNA based approaches. b, Rarefaction curve of sequencing reads at increasing depth. c, PCOA analysis of bacterial diversity across all samples using weighted UniFrac distance metric. d, Predictive metabolism profiling of RNA and DNA-based microbiomes of broiler ceca and intestine using PICRUSt at level 1. e, Culture based enumeration of different members of selected phyla using selective and/or differential media (Supporting information table 3). f, Ratio of total load of Firmicutes and Bacteroidota from DNA and RNA based amplicon sequencing data and culture-based (CB) analysis of ceca and intestinal contetns. g, Firmicutes:Bacteroidota ratio of ceca and intestine through DNA and RNA based microbiomes and culture-based (CB) approach

The predictive metabolism profiles (level 1) generated using PICRUSt also have variations in the DNA- and RNA-based microbiomes (Figure 2d). Here again, although predictive, the RNA-based microbiomes correctly represented the metabolism profiles (Supporting information Figure S4). For example, the carbohydrate metabolism and membrane transport are the active components in the intestine, whereas glycan biosynthesis and metabolism is active in the ceca.

## 4.0 Discussions

The first few works, dealing with the extraction of bacterial RNA relied mostly on ultracentrifugation (Kirby, 1964; Kirby, Fox-Carter, & Guest, 1967). In 1972, a simpler method, particularly targeting mRNA from eukaryotic tissues was developed (Brawerman, Mendecki, & Lee, 1972), which was later used for microbial RNA extraction by Gopalkrishna et al. (Gopalakrishna, Langley, & Sarkar, 1981). During this time frame, Birnboim and Doly developed their legendary method for the extraction of plasmid DNA from bacteria using the alkaline lysis method (Birnboim & Doly, 1979), which became a cornerstone for developing our RNA extraction protocol. The concept of eliminating genomic DNA and protein contamination, based on their denaturation followed by mis-folded renaturation was utilized in our RNA extraction protocol. Specifically, we used a variant of the alkaline lysis solution I, albeit with much lower pH of 4.7 adjusted using acetic acid (plasmid isolation use Tris of pH 8.4). The decision to use a low pH cell suspension solution was inspired by earlier works of RNA extraction methods (Marchiani & Aragno, 1995; Moran, Torsvik, Torsvik, & Hodson, 1993; Rivas et al., 2001), which otherwise are not suitable for diverse environmental samples and require a significantly larger number of reagents. The concept of using water-saturated phenol (pH 4.7) followed by heat treatment (95 °C for 4 minutes) for cell lysis was developed following earlier research (Nwokeoji, Kilby, Portwood, & Dickman, 2016; Sung, Khan, Nawaz, & Khan, 2003). But these methods either lysed cells at very high temperatures for longer periods (which led to RNA degradation) or used a silica membrane column, which is costly. The acidic pH helps stabilise RNA and inhibits RNase activity (Chheda et al., 2024; Kumar, Sharma, Kumari, Jagannadham, & Debnath, 2011), while thermal denaturation ensures the unfolding of DNA and protein structures. Subsequently, incubating the lysate in ice promotes erroneous renaturation of the denatured DNA and proteins. Incubation on ice decreases molecular motion, reducing the solubility of denatured proteins and DNA, and they aggregate as their thermal energy becomes insufficient to maintain solubility and form white precipitation. During the phenol-chloroform extraction step, these misfolded DNA-protein complexes partition into the interphase and organic phase due to their hydrophobicity and aggregation, while the intact RNA remains in the aqueous phase. This step is critical in ensuring selective recovery of RNA and effective removal of contaminating DNA and protein. The RNA finally suspended in autoclaved double-distilled water (pH 6.5) remained stable at −20 °C for 6 months (Figure 1f). RNA is stable at low pH because protonation reduces negative charge repulsion, leading to a less reactive 2’-hydroxyl group of ribose with minimised nucleophilic activity, preserving the phosphodiester bonds in the RNA backbone (Kornienko et al., 2024). The low pH of all the reagents used in this protocol ensured RNA stability (Figure 1b and 1f). For visualisation of RNA in agarose gel, we used 1X TAE buffer (pH 7.0) instead of the reducing MOPS-formaldehyde gel, commonly used for RNA visualisation. In the agarose gel, three distinct bands corresponding to 23S, 16S and 5S rRNA were observed with a higher intensity of the larger 23S rRNA band than the other two (Figure 1b). A common challenge in RNA extraction is the co-isolation of genomic DNA, often necessitating an additional DNase treatment to ensure RNA purity. In our method, we obtained DNA-free RNA as confirmed through agarose gel electrophoresis and negative PCR reaction of 16S rRNA gene, eliminating the need for post-extraction DNase treatment (Figure 1b and 1c).

Compared to commercial kits such as the Trizol reagent and Qiagen RNeasy Mini Kit, which are significantly more expensive and time-consuming (∼INR 150–500 per reaction; and ∼60-120 mins), our method offers similar or superior yield at a fraction of the cost (∼INR 5 per reaction; and ∼30 mins) (Table 1). The major reason being the absence of commonly used costly consumables, such as diethyl pyrocarbonate (DEPC), and RNase-free plasticware/water. Also, unlike many conventional RNA extraction protocols (discussed above), no chaotropic agents, detergents, or bead-beating steps are required, significantly reducing reagent costs and processing time (Table 2).

The first effort for the isolation of total RNA from complex microbiome samples, such as soil, was taken by Hahn et al. in 1990 (Hahn, Kester, Starrenburg, & Akkermans, 1990). Tsai et al. isolated mRNA directly from soil in 1991 (Tsai, Park, & Olson, 1991). Moran et al. developed a method for the isolation of rRNA from soil and sediment samples for ecological studies (Moran et al., 1993). But these methods were very critical, cumbersome and used a large number of reagents. Felske et al. developed an ultracentrifugation-based rRNA isolation method from soil and found the rRNA-based community to represent the metabolically active members (Felske, Engelen, Nübel, & Backhaus, 1996). Thorn et al. developed a cost-effective and robust phenol-chloroform-based protocol for the co-extraction of DNA, RNA, and proteins from complex microbiomes, including diverse soil types and found that RNA-based microbiome differs from DNA-based ones (Thorn et al., 2019). But, in this method, the use of DNase was necessary. Poursalavati et al. using bead beating, isolated RNA from soil and performed Oxford Nanopore sequencing for meta-transcriptomics analysis (Poursalavati, Javaran, Laforest-Lapointe, & Fall, 2023). There had been many similar meta-transcriptomics studies for gut environments (Bashiardes, Zilberman-Schapira, & Elinav, 2016; Gosalbes et al., 2011; Hatch et al., 2019; Ojala, Kankuri, & Kankainen, 2023; Schirmer et al., 2018). Even, many studies compared the gut metagenomic and meta-transcriptomics data and found significant differences in the distribution of bacterial members in both of these approaches (Franzosa et al., 2014; Jovel et al., 2022; Peters et al., 2019; Wang, Zhang, Liu, Zhang, & Sun, 2021; Yang et al., 2022). But, majority of these studies used expensive commercial kits for RNA isolation.

To evaluate the effectiveness of our newly developed meta-RNA isolation method, a comparative, 16S rRNA gene amplicon sequence analysis of the gut microbiome of broiler birds was conducted on the RNA-derived cDNA (active microbiome) and conventional metagenomic DNA (total microbiome). The use of two anatomically and functionally distinct regions of the broiler gut, i.e., the ceca and the intestine, was directed by their well-established physiological and environmental differences. The ceca, with its slower digesta transit, anaerobic conditions, and high fermentative activity, supports a rich community of obligate anaerobes and polysaccharide-degrading bacteria (Józefiak, Rutkowski, & Martin, 2004). In contrast, the intestine presents a more oxygenated, rapidly transiting environment that favours facultative anaerobes and fast-growing taxa (Yadav & Jha, 2019). These distinct ecological niches provide an ideal framework to evaluate the effectiveness of RNA-based microbiomes to profile the active microbial constituents. Since DNA-based microbiome profiles include both active and dead members, we initially hypothesised that they would exhibit a higher number of observed features (i.e., OTUs). However, contrary to our expectation, the RNA-based microbiomes from both the ceca and the intestine showed a greater number of OTUs. This unexpected result was possibly due to the fact that the RNA-based approach more accurately captures the transcriptionally active members of the community, including low-abundance taxa that may be under-represented or obscured in DNA-based analyses by the presence of relic DNA from dead cells. In the DNA-based approach, ceca were mostly populated by Firmicutes (78.57%), while the intestine by Proteobacteria (86.70%). But, in the RNA-based microbiome, although the major phyla remained the same, but their dominance got almost halved. While, the ceca had Firmicutes (52.53%) and Bacteroidota (31.45%) as the major members; the intestine had Proteobacteria (47.23%) and Firmicutes (44.03%) (Figure 2a). In the ceca, the DNA-based microbiome profile showed a clear over-representation of Firmicutes and a significant under-representation of Bacteroidota. In contrast, the RNA-based microbiome provided a more accurate and balanced depiction, with both Firmicutes and Bacteroidota being present in substantial proportions, which corroborates to the culture-based enumerations (Firmicutes and Bacteroidota load in the ceca were 1.6 × 10^12^ CFU g^-1^, and 1.7 × 10^12^ CFU g^-1^ respectively). This aligns with the known fermentative role of the ceca, where these two phyla are central to the breakdown of complex polysaccharides and the production of short-chain fatty acids (SCFAs) (Li et al., 2024; Wassie et al., 2022). Specifically, the RNA-based data identified Clostridia (Firmicutes) and Bacteroidia (Bacteroidota) as the dominant classes, both of which include genera known to specialise in anaerobic fermentation and fibre degradation (Chai et al., 2019; Döring & Basen, 2024; Zhang et al., 2022). The selective plating approach using reinforced clostridial agar (RCA) and bile esculin agar supports this observation. RCA, which favours the growth of *Clostridium*, yielded a higher count in the ceca (1.4 × 10^12^ CFU g^-1^) compared to the intestine (6.7 × 10^8^ CFU g^-1^). Similarly, *Bacteroides*, the major genus within Bacteroidota, was enumerated using bile esculin agar supplemented with vitamin K_1_, hemin, and kanamycin, resulting in 6.0 × 10^11^ CFU g^-1^ in the ceca. These results reinforce the conclusion that Clostridia and Bacteroidia are not only present but metabolically active in the cecal environment, contributing to its specialised fermentative functions. In contrast, the intestinal microbiome presents a different ecological niche. DNA-based sequencing data showed a pronounced overrepresentation of Proteobacteria, particularly the class Gammaproteobacteria. A similar observation, with more than 90% sequences representing Proteobacteria, was made by an earlier study (Wei, Morrison, & Yu, 2013). This could again be attributed to relic DNA from dead or lysed cells, which can persist in the intestinal contents and distort the community profile. However, the RNA-based microbiome depicted a more realistic representation of active taxa, revealing that Firmicutes and Proteobacteria were present in comparable proportions. Within Firmicutes, the Bacilli class was predominant, which includes well-known probiotic and commensal genera such as *Lactobacillus* and *Enterococcus*. These microbes are not only rapidly growing and metabolically versatile but also play critical roles in nutrient absorption, immune modulation, and antimicrobial production, making them essential contributors to intestinal health and function (Bazireh, Shariati, Azimzadeh Jamalkandi, Ahmadi, & Boroumand, 2020; Mountzouris et al., 2007). This RNA-based data was again validated by culture-based enumeration on MRS agar, selective for lactic acid bacteria, which revealed a higher Bacilli load in the intestine (6.3 × 10^10^ CFU g^-1^) compared to the ceca (1.2 × 10^10^ CFU g^-1^). Similarly, although the Bacteroidota phylum was less abundant in the intestine overall, both RNA-based sequencing and culture methods detected lower but consistent levels, with total Bacteroidota CFUs being 1.1 × 10^9^ g^-1^ and *Bacteroides* CFUs 6.8 × 10^8^ g^-1^.

The Firmicutes: Bacteroidota (F:B) ratio is considered a critical indicator of gut health, primarily because these two dominant bacterial phyla play contrasting but complementary roles in gut physiology and host metabolism (An, Kwon, & Kim, 2023; Burananat et al., 2025; Petakh, Oksenych, & Kamyshnyi, 2023; Stojanov, Berlec, & Štrukelj, 2020). Firmicutes are typically associated with the breakdown of complex carbohydrates and enhanced energy harvest, contributing to nutrient absorption and sometimes linked with weight gain and metabolic disorders. Bacteroidota, on the other hand, are known for their specialisation in polysaccharide degradation and production of short-chain fatty acids (SCFAs), which are essential for maintaining gut epithelial integrity, modulating immune responses, and ensuring metabolic stability (Lee & Lee, 2020; Zhao et al., 2025). Therefore, the balance between these phyla, the F:B ratio, serves as a functional marker of microbial homeostasis, metabolic activity, and the general health of the gut ecosystem. Interestingly, in our study, the F:B ratios obtained from the RNA-based analyses are in agreement with the culture-based data. While the ratios were 1.67 and 112.33 for ceca and intestine respectively; the same values from culture-based enumerations were 1.38 and 69.19, respectively. In sharp contrast, the DNA-based microbiome had an F:B ratio of 9.56 and 1.95 respectively in ceca and intestine (Figure 2f). Similar patterns were observed in our previous research involving poultry faecal samples, where the DNA-based microbiome showed a higher F:B ratio compared to culture-based results (Banerjee et al., 2018).

Based on our findings, we propose a new microbial ratio, Firmicutes to Proteobacteria (F:P), as an additional and potentially valuable indicator of gut health, particularly in the intestinal environment. While the Firmicutes: Bacteroidota (F:B) ratio has long been recognised for its relevance in microbial balance and metabolic function, it does not fully capture the dysbiotic shifts often marked by the overgrowth of opportunistic pathogens, many of which belong to the phylum Proteobacteria. Proteobacteria, especially members of the class Gammaproteobacteria, are frequently associated with inflammation, oxidative stress, and gut barrier dysfunction. A disproportionate increase in Proteobacteria is considered a hallmark of microbial dysbiosis and is linked to various gut-related diseases (Dam, Misra, & Banerjee, 2019; Shin, Whon, & Bae, 2015). The RNA microbiome-based F:P ratio was again found to align well with culture-based enumerations, in contrast to the DNA-based data (Figure 2g). Thus, the Firmicutes: Proteobacteria ratio offers a complementary perspective to the F:B ratio, by not only reflecting the contribution of beneficial, fermentative Firmicutes (such as Bacilli) but also capturing the overrepresentation of Proteobacteria, a sign of microbial imbalance and potential gut dysfunction. This ratio may prove especially useful in identifying early stages of intestinal dysbiosis, monitoring gut microbial resilience, and evaluating the impact of dietary, probiotic, or antimicrobial interventions in both poultry and broader gut microbiome research.

The predictive functional profiling further supports the accuracy of RNA-based microbiome analysis in deciphering the metabolically active microbial community. Using PICRUSt-based level 1 pathway predictions, distinct variations were observed between DNA- and RNA-based microbiomes (Figure 2d). In the RNA-based profiles, carbohydrate metabolism and membrane transport pathways were found to be predominant in the intestine, aligning with its physiological role in nutrient assimilation, rapid transit, and interaction with dietary substrates. Conversely, the ceca, a fermentation-rich, anaerobic chamber, exhibited high activity in glycan biosynthesis and metabolism, reflecting the breakdown of complex polysaccharides and synthesis of short-chain fatty acids by obligate anaerobes (Supporting information Figure 4) (Banerjee et al., 2018). These pathway predictions correspond well with the known ecological and metabolic functions of the respective gut regions and reinforce the value of RNA-based microbiome analysis as a more reliable tool for gut microbial ecology and health assessment.

## Conclusion

This study demonstrates the superiority of RNA-based microbiome analysis in capturing active microbial populations and their functional potential. By introducing a robust and economical RNA isolation protocol, the study paves the way for more accurate and accessible microbiome research. The findings underscore the importance of RNA-based methodologies in unravelling the functional dynamics of complex microbial communities, particularly in gut ecosystems.

## Data availability

All the raw sequences used in this study have been submitted to NCBI under the bioproject PRJNA1250238.

## Acknowledgement

We acknowledge DBT [BT/PR28574/AAQ/3/920/2018] and SERB [CRG/2021/007475] grants for providing financial assistance during the course of this study.

## Author contributions

A.C. and B.D. designed and conceived the study. B.D. Supervised the project. A.C. performed all the experiments, analysed the data and wrote the initial drafts. B.D. reviewed and edited the drafts.

## Competing interests

The authors declare no competing interests.

---

**Extended data table 1:** Comparison of the RNA extraction method (this study) with other commercially available kits. Data represented as mean ± standard deviation

**Extended data table 2:** Comparison of the developed RNA extraction process with other methods. A search with the ((TS=(RNA extraction)) AND TS=(Bacteria)) AND TS=(Method) was done in web of science. After screening 638 results, those were selected for comparison, where indigenous RNA extraction method was developed in last 25 years.

## Notes

### Competing Interest Statement

The authors have declared no competing interest.

