## Supplementary information for "A rapid, cost-effective, efficient method for total RNA extraction from pure bacterial cultures and complex microbiomes and its effectiveness in deciphering functionally active microbial populations"

**Article type:** Resource article

Running title: meta-RNA-based analysis of gut microbiomes

**Supporting information Table 1: Details of culture media used for enumeration of members of different phyla**

| Targeted phylum | Name of the media | Type of media | Growth parameters | | | | Target bacteria  (% Coverage) | Specific colony morphology of the target bacteria | Amount to mix in water (gm/1000ml) | Selective agent (Chromogenic substance) | References |
| --- | --- | --- | --- | --- | --- | --- | --- | --- | --- | --- | --- |
|  |  |  | **Temp (°C)** | **pH** | **O_2_** | **Time (Hrs)** |  |  |  |  |  |
| Firmicutes | Hicrome M-Enterococci agar^#^ | Selective, Differential | 37 | 7.8±0.2 | Yes | 18-24 | *Enterococcus faecalis* (50%) | blue | 27.1 | Cephalexinaztreonam (Phenol red and chromogenic substrate) | (Willinger & Manafi, 1995) |
|  |  |  |  |  |  |  | *Enterococcus faecium* (50%) | green |  |  |  |
|  |  |  |  |  |  |  | *Enterococcus hirae* (50%) | Blue |  |  |  |
|  | Lactobacillus MRS agar | Selective | 37 | 6.5 | No | 48 | *Lactobacillus* sp*.* (70%) | Large, round, creamy white, opaque colonies | 55.15 | Sodium acetate, Ammonium citrate (Absent) | (de Man, Rogosa, & Sharpe, 1960) |
|  | Mannitol salt agar | Selective, Differential | 30-35 | 7.2±0.2 | Yes | 18-72 | *Staphylococcus* sp. (70%) | yellow/white colonies surrounded by yellow zone. | 111.02 | Mannitol salt, Sodium Chloride  (Phenol Red) | (Chapman, 1945) |
|  | Bile esculin agar, (with kanamycin (20µg/ml) and sodium azide (0.15 mg/ml)) | Selective | 37 | 7 | Yes | 48 | *Lactococci and Streptococci* (80%) | Small, translucent colonies surrounded by a black halo | 64.5 | Kanamycin,  Sodium azide  (Esculin and FAC) | (Banerjee et al., 2018) |
|  | Reinforced clostridial agar | Enrichment | 35 - 37 | 6.8±0.2 | No | 24-48 | *Clostridium sporogenes*  *Clostridium perfringenes* | Irregular, finely scalloped, grayish and pitted; or smooth, flat, regular opaque, grayish, and pitted colonies | 38 | Sodium sulphite, Ferric citrate (Resazurin) | (Hirsch & Grinsted, 1954) |
| Bacteroidota (Previously Bacteroidetes) | HP6 agar (glucose (50mg/ml)) | Selective | 30-35 | 7.0±0.2 | Yes | 24 | *Cytophaga heparina* (50%) | Bluish green colony | 27 | Glucose (Absent) | (Christensen, 1977) |
|  | Nutrient agar (with kanamycin (50µg/ml)) | Selective | 37 | 7 | Yes | 24 | *Flavobacterium sp* (70%) | Yellow (cream to orange), round or tapered end colonies with prominent spread on agar surface due to gliding motility. | 28 | Kanamycin (Absent) | (Lapage, Shelton, & Mitchell, 1970) (Banerjee et al., 2018) |
|  | Bile esculin agar [with vit. K1 (0.01 mg ml-1), hemin (0.01mg ml-1), kanamycin (50 µg ml-1)] | Selective | 37 | 7 | No | 48 | *Bacteroides sp* (75%) | Gray, entire (unbroken), circular, raised colonies with blackish colouration in the medium | 64.5 | Kanamycin, Vit K1, Hemin (Esculin and FAC) | (Banerjee et al., 2018) |
| Proteobacteria | HiCrome *E. coli* Agar | Selective, Differential | 37 | 7.2±0.2 | Yes | 18-24 | *E. coli* (50%) | Bluish green colony | 36.57 | Bile salts (X-Glucuronide) | (Anderson & Baird‐Parker, 1975) |
|  |  |  |  |  |  |  | *Salmonella* Enteritidis (50%) | colourless |  |  |  |
|  | Hicrome *Klebsiella* Agar | Selective, Differential | 37 | 7.1±0.2 | Yes | 18-24 | *Klebsiella pneumoniae* (50%) | Purple-magenta (mucoid) | 20.04 | Bile salt, SLS, Carbenicillin (chromogenic mixture) | (Krieg & Holt, 1984) |
|  | Hicrome *Vibrio* Agar^#^ | Selective, Differential | 37 | 8.5±0.2 | Yes | 18-24 | *Vibrio cholerae* (50%) | Purple | 67.5 | Alkaline pH | (Hara-Kudo et al., 2001) |
|  |  |  |  |  |  |  | *Vibrio vulnificus* (50%) | Light purple to purple |  |  |  |
|  |  |  |  |  |  |  | *Vibrio parahaemolyticus* (50%) | Light green |  |  |  |
|  | *Salmonella-Shigella* Agar^#^ | Selective, Differential | 27 | 7.0±0.2 | Yes | 18-24 | *Salmonella typhi* (50%) | colourless with black center. | 63.02 | Bile salts, Brilliant Green, Sodium citrate (Neutral Red) | (Balows, Lennette, Hausler, & Shadomy, 1985) |
|  |  |  |  |  |  |  | *Shigella flexneri* (50%) | Small, colourless to redish colonies. |  |  |  |
| Campilobacterota | *Campylobacter* agar | Selective | 27 | 7.4±0.2 | No | 24 | *Campylobacter sp* | Small, Yellowish to grey, round, convex, glistening surface, Translucent colonies. | 45 | Hemin,Vancomycin, Ammphotericin B, Cycloheximide, Cefoperazone | (Manning et al., 2001) |

**#** Denotes non autoclavable media

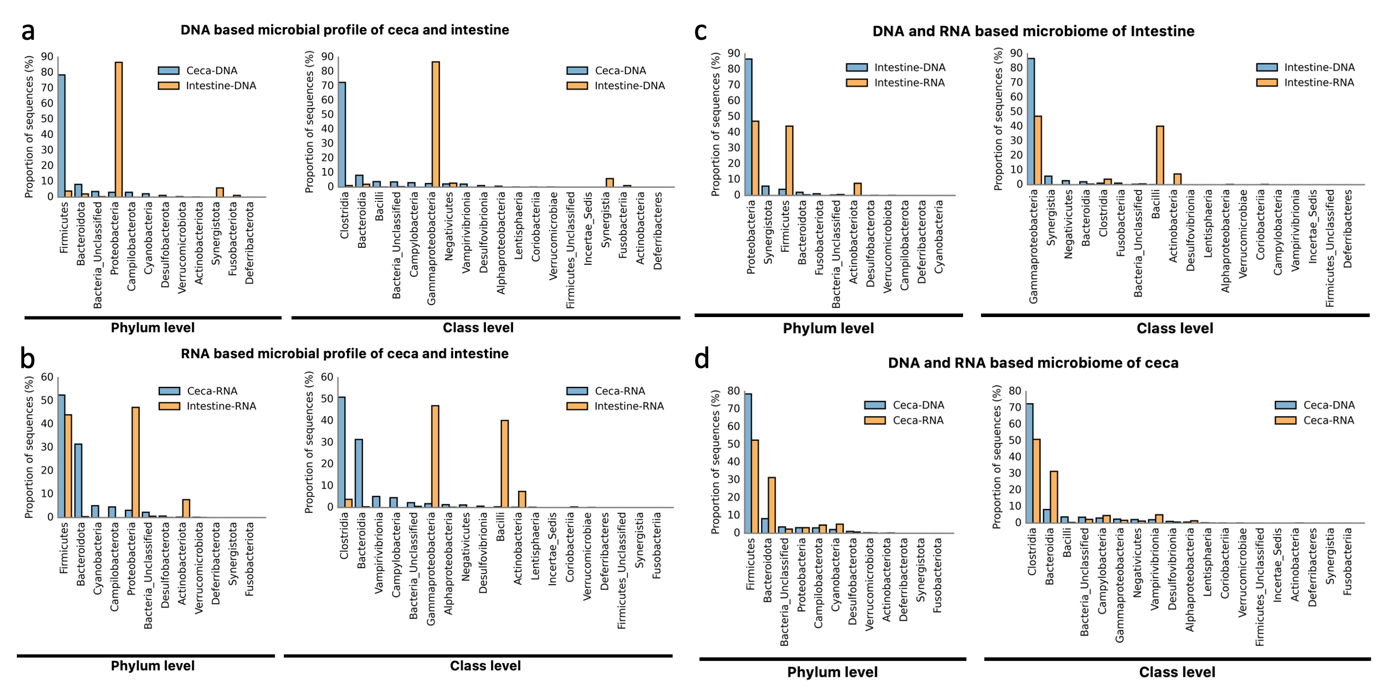

**Supporting information Figure S1: Distribution of different phyla and classes across ceca and intestine.** a, Relative proportions of bacterial members in the DNA based amplicon data of ceca and intestine at phylum and class level b, Relative distribution of phylum and class in ceca and intestine in RNA-based microbiome. c, RNA-based microbiome of ceca is different from DNA based analysis. d, RNA-based microbiome of intestine is different from DNA-based microbiome

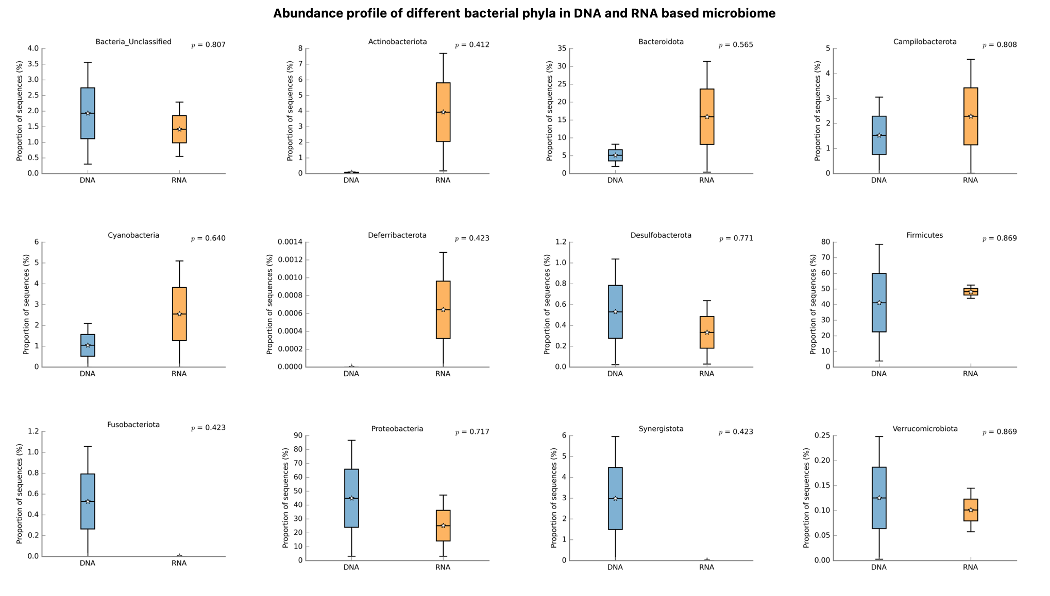

**Supporting information Figure S2: Distribution of different phyla in DNA and RNA based microbiome.** For the calculation of distribution of different phyla across DNA and RNA-based microbiomes, both samples of ceca and intestine were combined and calculated as single samples for DNA and RNA

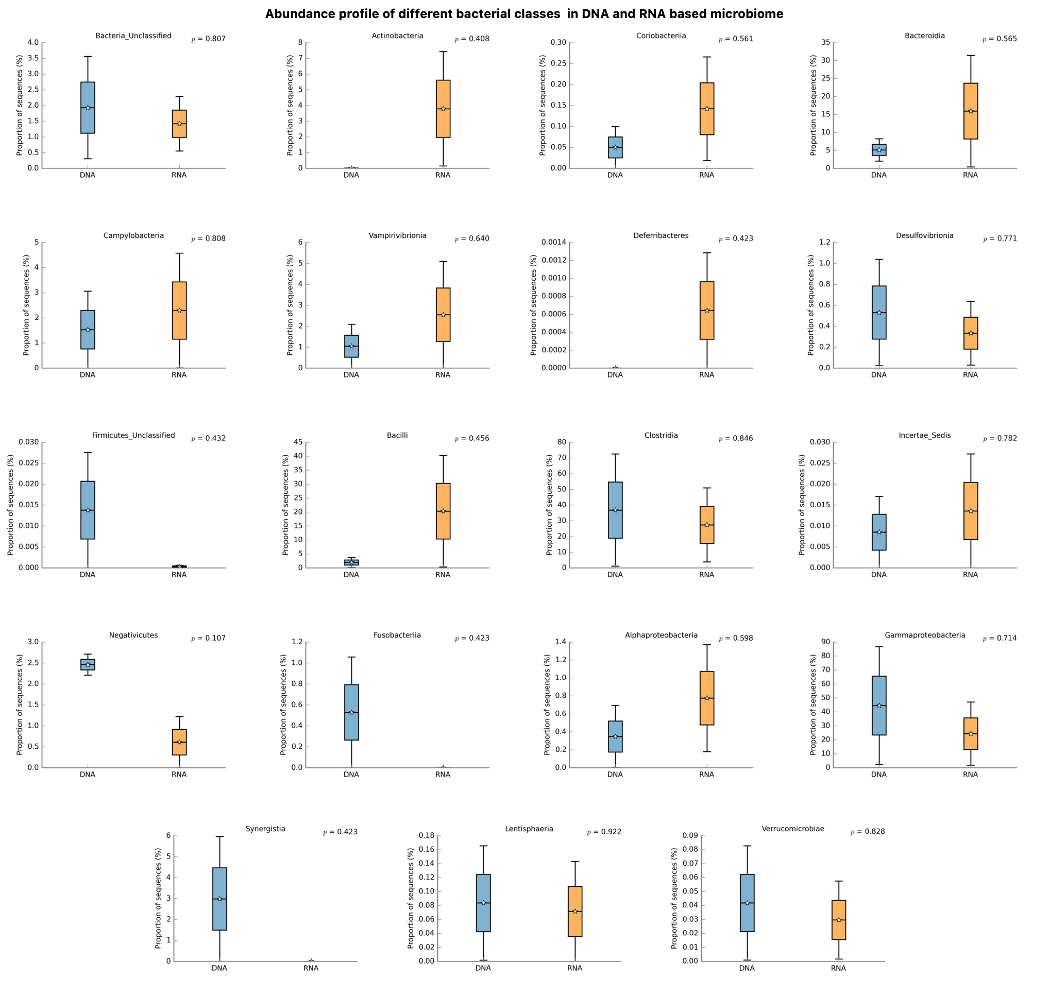

**Supporting information Figure S3: Distribution of different classes in DNA and RNA-based microbiome**. For the calculation of the distribution of different phyla across DNA and RNA-based microbiomes, both samples of the ceca and the intestine were combined and calculated as a single sample for DNA and RNA

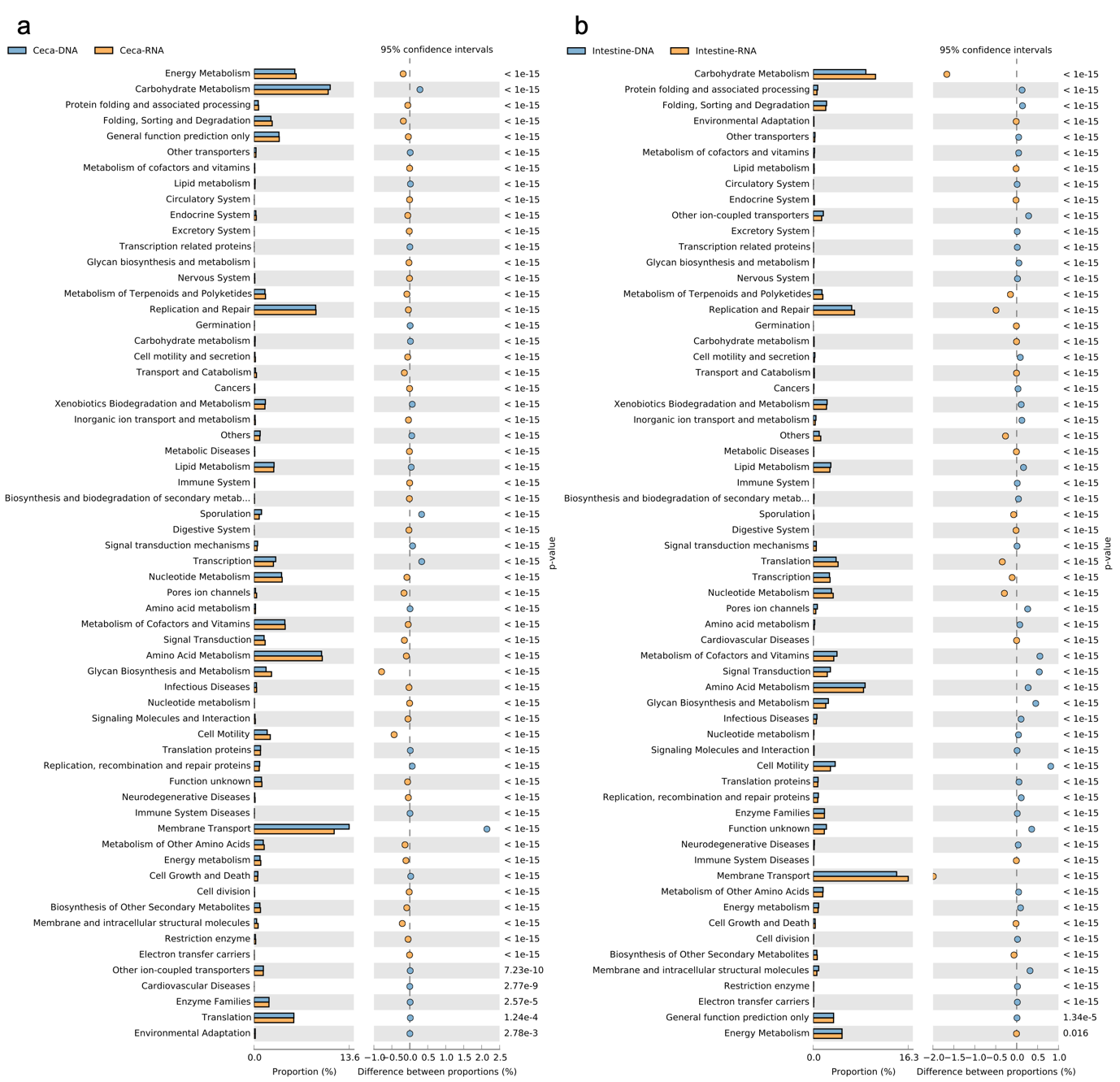

**Supporting information Figure S4: Predictive metabolism of ceca (a) and intestine (b) in DNA and RNA-based microbiomes using PICRUSt at level 2**

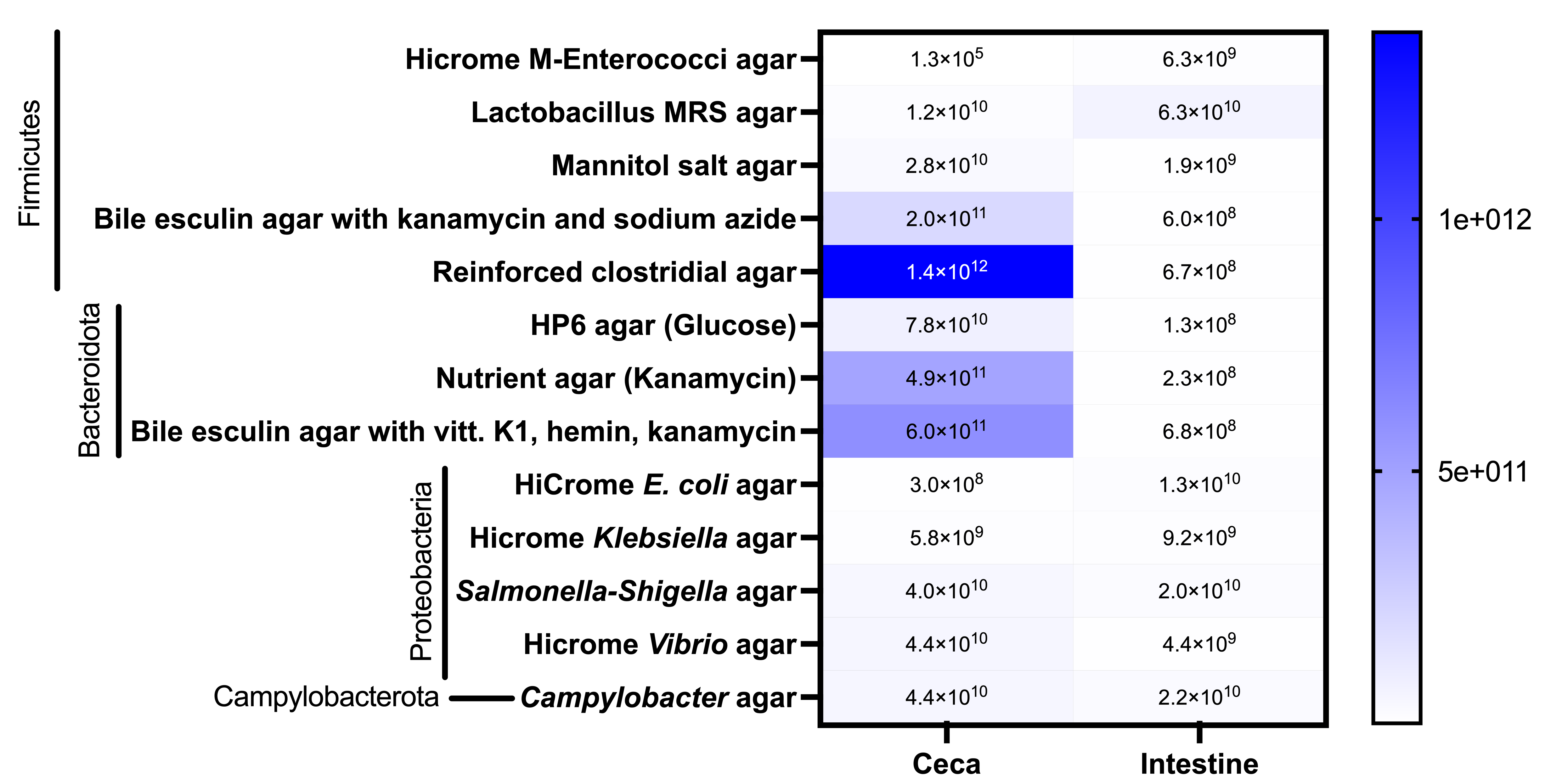

**Supporting information Figure S5: Culture-based enumeration of members of different bacterial phyla in ceca and intestine of broiler birds**
